# Proteochemometric method for pIC50 prediction of Flaviviridae

**DOI:** 10.1101/2022.03.16.484682

**Authors:** Divye Singh, Avani Mahadik, Shraddha Surana, Pooja Arora

## Abstract

Viruses remain an area of concern despite constant development of antiviral drugs and therapies. One of the contributors among others is the flaviviridae family of viruses. Like other spaces, antiviral peptides (AVP) are gaining importance for studying flaviviridae family. Along with antiviral properties of peptides, information about bioactivity takes it even closer to accurate predictions of peptide capabilities. Experimental identification of bioactivity of each potential peptide is an expensive and time consuming task. Computational methods like Proteochemometric modelling (PCM) are promising for prediction of bioactivity based on peptide and target sequence. The additional edge PCM methods bring in is the aspect of considering both peptide and target properties instead of only looking at peptide properties. In this study, we propose prediction of pIC50 for AVP against flaviviridae family target proteins. The target proteins were manually curated from literature. Here we utilize the PCM descriptors as peptide descriptors, target descriptors and cross term descriptors. We observe taking peptide and target information improves the results qualitatively and gives better pIC50 predictions. The R^2^ and MAPE values are 0.85 and 8.44 % respectively

## I. Introduction

Viral diseases have been a cause of multiple epidemic outbreaks in last few decades. This includes may different viruses like Ebola, Zika, Dengue, SARS and other viruses. Although belonging to one bigger group of viruses they differ a lot in their activity, sequence, structure and function. Viruses are also known to have continuous mutations which makes it necessary and complex to identify antiviral drug candidates. This leads to the need for continuous drug development. Recently peptide-based antivirals have gained a lot of importance and have shown promising development[1]. Peptide-based reagents are advantageous due to their specificity, safety, suitability, and flexibility[2].

In this work, methods to predict the pIC50 value for Flaviviridae family are studied. Most common Flaviviridae viruses are dengue virus (DENV), Zika virus (ZIKV), yellow fever virus (YFV), Japanese encephalitis virus (JEV), West Nile virus (WNV), tick-borne encephalitis virus (TBEV), and hepatitis C virus (HCV), with the publically available antiviral peptides having maximum target proteins of HCV and DENV. Rajput et al. developed an algorithm to identify inhibition activity of chemicals from ChEMBL and peptides from AVP-pred databases against Flaviviruses using QSAR method [3]. Recently, Geoffrey et al. developed machine learning based Auto-QSAR using PubChem data, which generated drug leads for Flaviviruses. Further in silico modelling was performed for the drug leads and their target proteins [4].

Generally, antiviral peptides are studied based on physicochemical properties, evolutionary properties and profiles based on only peptide attributes[5]. However, a peptide can have good physicochemical properties that are similar to other bioactive peptides but its efficacy cannot be identified unless its bioactivity is experimentally determined. The Inhibition constant (IC) 50 value is commonly used for validating the activity of a peptide. Experimental determination of IC50 is a expensive and tedious task. Taking all the potential peptides for experimental validation might not be feasible. Prediction of IC50 values can reduce the time and effort and help in taking most promising peptide for further experimentation. There is very little research done in the area of in silico methods for IC50 value prediction. To understand the IC50 of a peptide it is necessary to understand the interaction with target protein. One of the method for doing so is proteochemometric modelling.

PCM is a computational method that can predict the bioactivity relations between ligand and targets. It is a method to incorporate the target interaction in sequence based analysis. Three types of descriptors are included in PCM-Target descriptor (captures information of target), Ligand descriptor (captures information of ligand), Cross-term descriptor (captures interaction between the ligand and its target). With the different types of interactions studied, scope of PCM has expanded to protein-peptide, protein-DNA and protein-protein interactions.

### A. Literature Survey

Recently Parks et al using ChEMBL25 dataset, generated proteochemometric models to predict pIC50 using random forest and feed-forward neural network [6]. The study checked the usability of PCM model to classify binders and nonbinders. For the ChEMBL25 data set, various physicochemical properties like log P, molecular weight, number of specific bonds and fingerprints were used as descriptors. Ventsislav Yordanov et al. demonstrated use of PCM for analysing structure-affinity relationship of antigen peptides binding to HLA-DP proteins [7]. The HLA system is an important part in the immune system. The HLA proteins bind to a wide range of antigenic peptides, which is essential for the immune recognition of the antigens. The chemical structure of peptides and proteins used, were described by three z-scales. Bio-activity modelling of multiple compounds against protein isoforms was done by Rasti et al. using proteochemometrics modeling [8]. They applied PCM to investigate inhibition of Carbon Anhydrase isoforms using combination of different descriptors (three zscale, five z-scale and GRIND) was used. Mutations affect the antimicrobial activity, PCM model have also been used to identify the mutations. The study by Nabu et al. helped in understanding the impact of physicochemical properties of mutated amino acids on resistance of penicillin binding proteins [9]. The mutation positions and various chemical descriptors were utilised as protein sequence and ligand descriptors respectively.

### B. Approach

In this work, a PCM based model for prediction of pIC50 values for peptides against flaviviridae family is developed. The peptides and target protein were cumulatively studied using PCM descriptors which includes peptide properties, z-scales for peptides, protein and peptide-protein interaction. The target proteins here were manually curated from literature.

The overall approach of the study includes:

- Curation of dataset
- Defining PCM descriptors
- Training on a suitable machine learning model.
- pIC50 prediction algorithm

## II. Materials and methods

This section elaborates on data, features and details of the machine learning algorithms.

### A. Curating the dataset

The datasets were created based on the publically available anti-viral peptide data with their IC50 values and the target proteins collected from literature. This resulted in two datasets:

- Antiviral peptides with IC50 values (Dataset 1) and
- Antiviral peptides with IC50 values and their target proteins (Dataset 2).

1. *Dataset 1:* Anti-viral peptide data was taken from publicly available AVP-IC50 dataset [10]. The dataset consists of AVP sequences, IC50 values in micromolar and their respective viral families. From these, peptides for only Flaviviridae family were taken constituting dataset 1 with total of 130 sequences.
2. *Dataset 2:* For dataset 2, alongwith the AVP, flaviviridae target proteins were also taken. The target proteins of the antiviral peptides were identified from literature [11]–[31]. 50 peptides sequences out of 130 had defined target. These 50 peptides sequences along with their target protein sequences formed dataset 2. Target protein sequences were extracted from Uniprot [32].
3. *pIC50:* The IC50 values in the datasets ranged from 0.001 micromolar to 440 micromolar making the distribution very skewed, as shown in 1(a). This would have made it difficult for the model to extract informative patterns to learn on. Thus, the IC50 values were negative log transformed to give pIC50 value. This is done as below formula

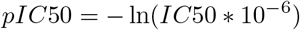

The data distribution after converting to pIC50 is shown in 1(b). Hereafter, pIC50 will be used in the rest of the paper.

**Fig. 1:**
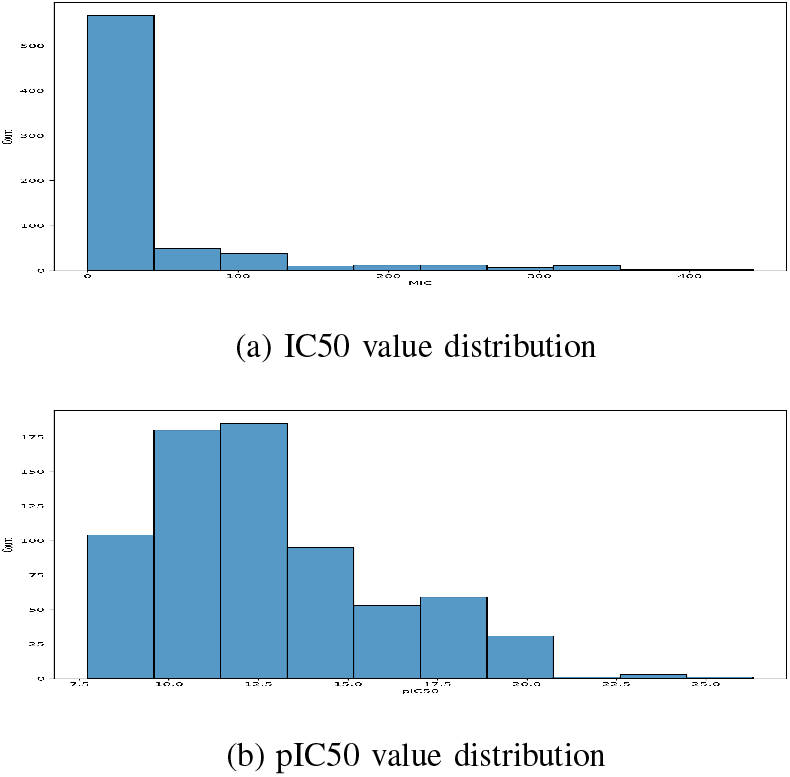
Distribution of IC50 and pIC50 values

### B. Descriptors

PCM modelling works based on descriptors which are mathematical representation of various properties of peptides and their target proteins. Here we are looking at the following descriptors which are calculated for peptides and their target proteins.

- Physicochemical properties of peptides and target protein
- Z-scale for peptides, target protein and their cross term.

#### Physicochemical Properties

Peptide and target protein properties were calculated from their amino acid sequences using Biopython package [33].

#### z-scales

The peptides and target proteins used in this work are represented using five z-scale descriptors (*z*_1_, *z*_2_, *z*_3_, *z*_4_ and *z*_5_) of their amino acid as derived by Sandberg et al. [34]. The z-scale represents their hydrophobicity (*z*_1_), steric bulk properties and polarizability (*z*_2_), polarity (*z*_3_), and electronic effects (*z*_4_ and *z*_5_). These five z-scales are the principal components of 26 computed and measured physicochemical properties of amino acids. To get a z-scale descriptor for a peptide and protein, the average is taken of their amino acid z-scale vectors. The z-scale descriptors for both peptides and proteins are normalised to standard normal. In order to incorporate the information about the interaction between protein and the peptide, cross-term descriptors were also included. This is calculated as flattened out outer product of normalised peptide z-scales and normalised protein z-scales resulting in 25 (5×5) dimensional vector. As a result, each peptide-protein pair is represented as 35 dimensional vector (5 peptide z-scale, 5 protein z-scale and 25 cross-term).

#### Descriptor groups

In order to perform various machine learning experiments, multiple groups of descriptors were created as defined below.

- Physicochemical properties for peptides (PP)
- Physicochemical properties for target protein (TP)
- Peptide z-scale descriptors (PZ) - 5 z-scale descriptors calculated for peptide sequences,
- Target protein z-scale descriptors (TZ) - 5 z-scale descriptors calculated for target protein sequences,
- Cross term descriptors (XZ) - Multiplication of peptide and target protein z-scale descriptors generated the cross term descriptors group.

### C. Machine learning details

In this section, we discuss the features selection method, machine learning algorithm and evaluation criteria of the model.

#### Feature selection

In machine learning, it is important to have useful input features or descriptors. Therefore, in-order to remove uninformative predictors, feature selection is carried out. This not only removes noise from the data, it also reduces dimensionality of the data which makes the trained model less complex and more interpretable. The feature selection for this work is carried out in two steps. First, the feature ranking is obtained using Recursive Feature Elimination. This is followed by adding the features iteratively starting from the highest ranking feature and checking for their predictive performance. Further, only those features were added to the final feature set whose addition improved the adjusted-*R*^2^ value. The calculation of adjusted-*R*^2^ based on *R*^2^ value is as

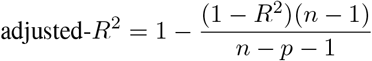

where, *R*^2^ is sample R-squared value, *n* is number of examples and *p* is number of predictors.

#### Random Forest

Random Forest is an ensemble learning method for classification and regression. The underlying principle is to construct multiple decision trees and aggregate the output from each decision tree. In case of regression problem, most common means of aggregating the output is mean of the predictions. In random forest, each of the decision tree is trained on randomly selected subset of features and examples from the training data which causes each tree to learn different patterns from the same training data. This results in a reduced variance, making the model more effective. Random forest regressor model from scikit-learn library[35] was used.

#### Evaluation criteria

Selection of informative performance metrics is vital in order to measure effectiveness of a prediction model. Therefore, to informatively measure performance of the prediction models, performance measures used in this work is described below:

- Root Mean Squared Error (RMSE):

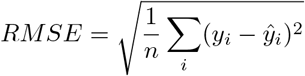
- Mean Absolute Percentage Error (MAPE):

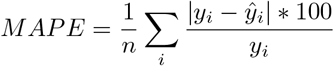
- R-squared value (*R*^2^):

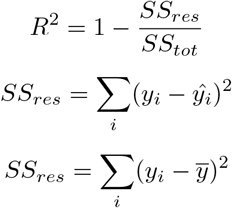

where, *SS_res_* is residual sum of squares, *SS_tot_* is total sum of squares
- Pearson’s Correlation Coefficient (PCC):

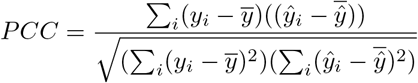

y is actual pIC50 value, 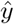 is predicted pIC50 value, 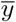 is mean of actual pIC50 values and 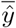 is mean of predicted pIC50 values.

### D. Methodology

This section elaborates on utilizing the curated data, properties and algorithm for training the suitable model. As explained in II-A properties were calculated for dataset 1 and dataset 2. Dataset 1 was studied using physicochemical properties. However, for dataset 2 all descriptor groups (as explained in section II-B) were calculated as it contains both peptide as well as target protein information. Using only the peptide properties for dataset 1 to train the model, resulted in lower *R*^2^ value. Considering this, further this study focuses on utilizing dataset 2.

We see an improvement in the results, with the peptide and target property groups included in the model. Using multiple sequence descriptors can be helpful for the model to learn. Hence, along with properties, z-scales were used to represent the sequences. The model was trained on peptide and target protein z-scales(PZ, TZ) which further improved the performance of the model. The PCM approach includes one more cross term group(XZ). A combination of property groups (PP, TP) and z-scale groups (PZ, TZ, XZ) was also tried, which performed the best of all the combinations. The descriptors calculated and the combinations tried are illustrated in 2. Further in order to remove the noisy or non-informative features, feature selection was carried out as explained in subsection II-C. During feature selection phase, Random Forest with default parameters was used as a base regression model.

The next step was to tune the Random Forest model on the selected features to get the best performance. For this, grid search approach was used where hyper-parameters form the axis of the grid and each point on the grid is a combination of defined values for each hyper-parameter. 5 fold cross-validation on Mean Absolute Percentage Error (MAPE) was used to obtain best performing hyper-parameter values. The hyper-parameters and their values for which the Random Forest model was tuned are mention in Table I. As a last step, the Random Forest was trained on the best preforming feature set and hyper-parameters and its predictive performance was evaluated. Further, to ensure fairness of the results all performance measures are calculated using Leave-one-out (LOO) cross-validation (CV).

**Fig. 2:**
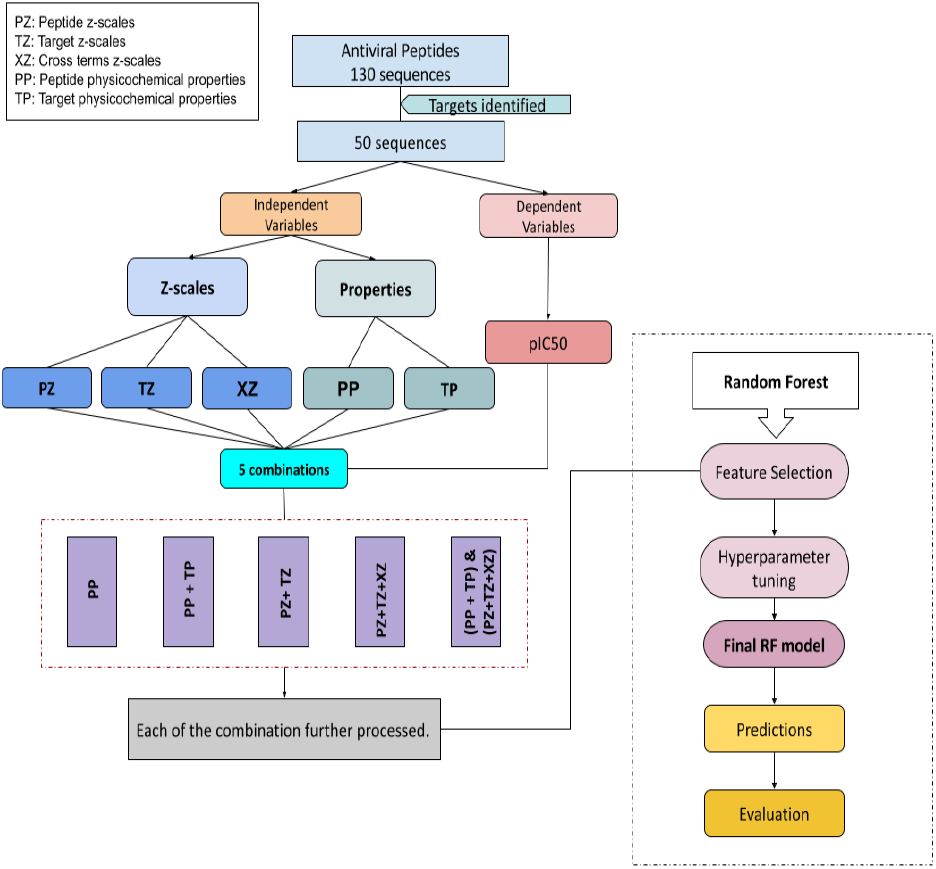
PCM model Flowchart: Illustration of the descriptors and multiple combinations of descriptor groups used to predict the pIC50 using Random forest.

**TABLE I:** Range of values used to tune hyperparameters for random forest

| Hyperparameter | Ranges |
| --- | --- |
| n_estimators | 100, 200, 250, 300, 500, 1000, **1500** |
| min_samples_split | **2**, 5, 10 |
| min_samples_leaf | 1, 2, **4** |
| max_features | **auto**, sqrt, log2 |
| max_depth | **2**, 3, 5, 10, 15, 20, None |
| bootstrap | True, **False** |

## III. Results and discussion

In this section we will look at the results obtained for various combinations of descriptor groups. Results for some of the example sequences is mentioned in Table II. It can be seen from the results that the predicted pIC50 values are very close to actual pIC50 values. The performance of the model was evaluated using Leave-one-out cross validation and the results are tabulated in Table III.

**TABLE II:**
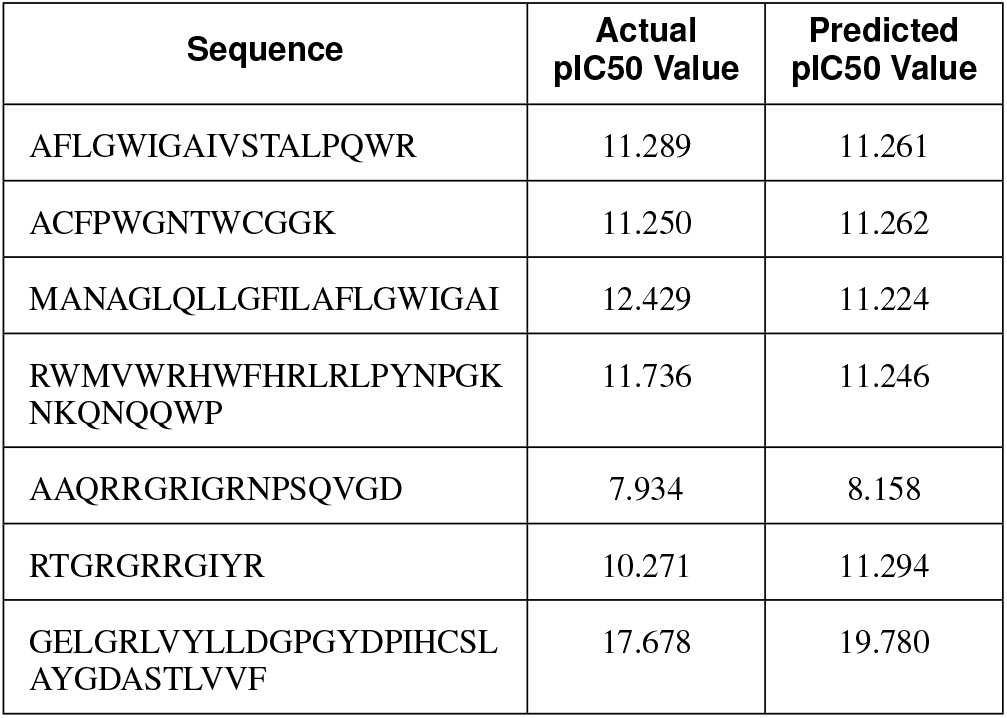
Actual and predicted pIC50 values for example sequences

**TABLE III:** Results of obtained models

| Descriptor combinations | R<sup>2</sup> | MAPE | PCC | MSE |
| --- | --- | --- | --- | --- |
| PP | 0.48 | 14.04 | 0.72 | 6.88 |
| PP + TP | 0.72 | 11.46 | 0.84 | 3.70 |
| PZ + TZ | 0.72 | 11.32 | 0.85 | 3.63 |
| PZ + TZ + XZ | 0.76 | 9.48 | 0.87 | 3.14 |
| (PP + TP) & (PZ + TZ + XZ) | **0.85** | **8.44** | **0.92** | **1.99** |

The model was trained using only peptides properties which resulted in the low r-squared value of 0.29 and high mean absolute percentage error of 17.10%. To understand this the dataset was analysed further. There are few peptide sequences in the dataset which are very similar but still have very different pIC50 value and vice verse. Since the physicochemical properties used were sequence based, the same was seen in property based descriptors. This might have it difficult for the model to learn informative patterns leading to poor results.

Based on the above results it was decided to incorporate information of target protein along with the peptides. As in peptide-protein interactions, not only the peptide but also the target on which the peptide acts is important, we tried incorporating target protein information in the model. Hence, PCM modelling approach, wherein target properties along with the peptide properties were also included was used. The adding of target protein properties improved the results in r-squared value of 0.30 and MAPE of 18.82%.

As described in section II-B, along with peptide properties Zscales were also calculated for each of the sequences which incorporate additional physicochemical properties of the peptides. This combination of Zscale for peptide and protein significantly improved the results giving r-squared value of 0.72 and MAPE of 11.32%.

However in PCM, along with the target and peptide descriptor groups, cross-term z-scales are often considered. The model was trained using the three descriptor groups (PZ, TZ and XZ). Addition of peptide–protein cross-term descriptor slightly improved the predictive performance of the model further, giving r-squared value of 0.76 and MAPE of 9.48%.

The combination of various descriptors can be helpful for the model, so we combined the z-scales and physicochemical properties. The combination resulted in further improved results with r-squared value of 0.85 and MAPE of 8.44%. This was the best result for all our experiments using different descriptor groups.

### A. Discussion

Despite various scientific discoveries and advancements viruses continue to be one of the threats to human health[36]. Antiviral peptides(AVP) are emerging as one of the interesting alternative therapeutics to viral concerns. Although elusive, antiviral peptides do exhibit certain physicochemical properties which makes them great candidates for antiviral therapeutics [37]. The work done by Surana et al. explained the usage of physicochemical properties[38]. Although physicochemical properties do help in identifying the potential antiviral peptides, IC50 is one of the methods used for further validating the efficacy of candidate peptides [39]. In order to predict the candidate peptides as close to the experimental method we propose the prediction of IC50 in our current study.

We initially utilized a combination of peptide properties and IC50 of known peptides to create the algorithm. However, we observed that the relationship between the peptides and the IC50 values cannot be directly correlated. The behavior of a peptide depends on its own properties as well as the nature of the target against which the peptide is going to act. This is where PCM methods seem to be the right approach where it does not only take only the peptides but also takes target protein into account.

PCM models give the flexibility to study multiple ligand(peptide in this case) and multiple target setup which becomes beneficial in the current study. As, for flaviviridae AVP although they are all against flaviviruses but have different targets that they act upon. As explained earlier the major features utilized were the physicochemical properties for peptides and target proteins. We observed that adding the physicochemical properties of target proteins improved the IC50 prediction for the peptides. Along with the basic physicochemical properties the addition of Z scores gave a boost to the model performance.

The Z score includes the hydrophobicity, steric bulk properties, polarity, polarizability and electronic effects giving additional features to the physicochemical properties. The Z scores individually covers the features of peptides, target proteins and cross descriptors covering the interaction features between the peptide and target proteins. The cross terms features give an additional edge in terms of covering the ligand - target interactions instead of only features of individual peptides and targets. Multiple combinations of physicochemical properties and Z-score descriptors lead us to the model with best descriptors giving good predictions. The best descriptors identified were target polarity, cross term of peptide polarity and target hydrophobicity along with aromaticity. It can be inferred that aromaticity along with hydrophobicity and polarity of protein and peptide best explain the relationship and hence the prediction of IC50.

We further explored PCM2Vec methodology for IC50 predictions. However, we did not get very good results there and need to be explored further. PCM2Vec can be a promising algorithm which can be explored in future work.

## IV. Conclusion

This study aimed to create a PCM model for predicting the pIC50 value for flaviviridae family. The IC50 prediction of probable antiviral peptide brings in an edge to give another layer of validation in terms of the peptide efficacy. As most of the peptides or drug candidates are build against specific targets, being able to predict their IC50 values is one step closer to getting more accurate peptides for further experimental validation and thus reducing the time and cost for experimental validation cycle. Here we propose an initial PCM method for AVP prediction. This can be further extended to multiple viral families or target based PCM model. Currently we have considered entire target sequences. Further experimentation can be done to include binding site residues and their interactions. Taking binding site residues into consideration might further boost the algorithm performance.

## V. Code and Data availability

The code and data is available at https://github.com/thoughtworks/mic-predictor

## VI. acknowledgement

We would like to acknowledge Dr. VK Jayaraman for their support and guidance.

We would like to thank Dr. Deepti Sahasrabuddhe for reviewing the paper.

IEEE conference templates contain guidance text for composing and formatting conference papers. Please ensure that all template text is removed from your conference paper prior to submission to the conference. Failure to remove the template text from your paper may result in your paper not being published.

